# Role of UCP2 in the energy metabolism of the cancer cell line A549

**DOI:** 10.1101/2022.09.13.507778

**Authors:** Jessica Segalés, Carlos Sánchez-Martín, Aleida Pujol, Marta Martín-Ruiz, Eduardo Rial

**Author notes:** **Corresponding author** Department of Structural and Chemical Biology, Centro de Investigaciones Biológicas Margarita Salas, CSIC, Ramiro de Maeztu 9, 28040 Madrid, Spain. Department of Experimental & Health Sciences, University Pompeu Fabra, Barcelona, Spain (Jessica Segalés). Department of Biosciences, University of Bari, Italy (Carlos Sánchez-Martín).

## Abstract

The uncoupling protein UCP2 is a mitochondrial carrier whose transport activity remains controversial. The physiological contexts in which UCP2 is expressed have led to the assumption that, like UCP1, it uncouples oxidative phosphorylation and as a result it lowers the generation of reactive oxygen species. Other reports have involved UCP2 in the Warburg effect and results showing that UCP2 catalyzes the export of matrix C4 metabolites to facilitate glutamine utilization, suggests that the carrier could be involved in the metabolic adaptations required for cell proliferation. We have examined the role of UCP2 in the energy metabolism of the lung adenocarcinoma cell line A549 and show that UCP2 silencing decreased the basal rate of respiration although this inhibition was not compensated by an increase in glycolysis. Silencing did not lead to changes in proton leakage, as determined from the rate of respiration in the absence of ATP synthesis, or changes in the rate of formation of reactive oxygen species. The decrease in energy metabolism did not alter the cellular energy charge. The decreased cell proliferation observed in UCP2-silenced cells would explain the decreased cellular ATP demand. We conclude that UCP2 does not operate as an uncoupling protein while our results are consistent with its activity as a C4-metabolite carrier involved in the metabolic adaptations of proliferating cells.

**Highlights:**

- UCP2 silencing decreases respiration without a compensatory increase in aerobic glycolysis
- ATP levels remain unchanged despite the reduction in energy metabolism
- UCP2 silencing decreases cell proliferation that could explain the decrease in energy demand
- UCP2 silencing does not change the proton leakage rate
- Data support the proposed involvement of UCP2 in the Warburg effect

## 1. Introduction

The uncoupling proteins (UCPs) are mitochondrial carriers which belong to the SLC25 family of transport proteins [1]. The name “uncoupling protein” should imply that the biochemical activity of these carriers should lower the efficiency of oxidative phosphorylation (OXPHOS). The uncoupling activity of UCP1 is well established and provides the molecular basis for heat production in brown adipose tissue [2]. Although UCP2 (SLC25A8) is a carrier that was initially named as an “uncoupling protein” due to its sequence homology to UCP1 [3], its transport activity and physiological roles are still under debate [4-12]. UCP2 is expressed in a variety of cells types and its expression levels have often been related to the defense against oxidative stress. Thus, uncoupling OXPHOS should, in principle, lead to an increase in the rate of respiration that should result in a lower generation reactive oxygen species (ROS) at the mitochondrial respiratory chain. However, the uncoupling activity of UCP2 and its role in the control of ROS formation remain as controversial issues [4,5,7,8,11,12].

The upregulation of UCP2 in tumor cells has raised considerable attention [13-19]. Besides its possible role in the control of ROS formation and chemoresistence, it has been proposed that UCP2 could also be of relevance for the metabolic reprogramming of the cancer cell. UCP2 has been involved in two events related to the Warburg effect: the use of glutamine as an energetic fuel and the export of Krebs cycle C4-metabolites out of mitochondria to be used in the biosynthesis of macromolecules [6,9,10]. This UCP2-mediated export of C4-metabolites has been proposed to also decrease the redox pressure on the respiratory chain, lowering ROS formation [9].

Here we evaluate the effect UCP2 silencing in the energy metabolism of the human lung adenocarcinoma cell line A549. We report that silencing lowers respiration while state 4 rates (respiration in the absence of ATP synthesis) remain unchanged. The decrease in OXPHOS is not accompanied by a compensatory increase in glycolysis while the cellular energy charge is maintained. These combined effects reflect a lower cellular demand for ATP that can be explained by the observed decrease in cell proliferation. We conclude that UCP2 does not present an uncoupling activity while our data support its involvement in the Warburg effect.

## 2. Materials and Methods

### 2.1. Cell culture and materials

Experiments were performed with the cell line A549 (human lung adenocarcinoma) obtained from the American Type Culture Collection (Manassas, USA). Cells were cultured at 37°C in a humidified 5% CO_2_ atmosphere in standard growth medium: DMEM supplemented with 10% heat-inactivated fetal bovine serum (FBS), 2 mM glutamine, and 100 U/mL of penicillin/streptomycin. All medium components were from Gibco (Life Technologies, Paisley, UK). Culture medium was changed every two days. Cell number and viability were assessed by Trypan Blue (0.2%) staining using a TC10 automated cell counter (Bio-Rad Laboratories, Hercules, California, USA). Reagents were from Sigma-Aldrich Merck (Darmstadt, Germany) unless otherwise stated.

### 2.2. Silencing

The day before transfection, A549 cells were seeded at a density that led to 50% confluence the following day (e.g., 12k cells/well for XF24 plates, 40k for p24 plates and 80k cells for p60 plates). A validated siRNA against the human UCP2 obtained from Ambion (s14630, Thermo Fisher Scientific, Lafayette, USA) and a negative scrambled control were used to transfect A549 cells using Lipofectamine 2000 (Invitrogen, Thermo Fisher Scientific, Lafayette, USA) following manufacturer’s protocol. RNAs (1 μM final) and lipofectamine were diluted in Opti-MEM, mixed 1:1 and incubated for 20 min at room temperature. Cells, in transfection medium (DMEM supplemented with 10% FBS and 2 mM glutamine without antibiotics), were transfected adding dropwise the siRNA/lipofectamine mix. Final siRNA concentration was 20 nM. The following day, transfection medium was replaced by standard growth medium and experiments carried out 24-48 hours later depending on the conditions.

### 2.3. Western blot analysis

UCP2 expression levels were determined by Western blot analysis as previously described [20]. 40 μg of cellular extracts were resolved by SDS-PAGE, transferred to nitrocellulose membranes and probed with an anti-UCP2 antibody (Ref. SC-6525, Santa Cruz Biotechnology, Santa Cruz, USA). Immunoblots were developed with the Super Signal West Dura chemiluminescent substrate (Pierce, ThermoScientific, Rockford, IL, USA).

### 2.4. Lactate assay

The rate of lactate formation in the culture media were determined enzymatically. Just before the assay, growth medium was replaced with fresh medium supplemented with FBS dialyzed against PBS to remove contaminating lactate. Duplicate samples collected 60 min later. Lactate concentration was determined with a colorimetric assay that follows NADH formation. Reaction buffer contained 1M glycine, 400 mM hydrazine, 20 mM EDTA pH 9.8. Reaction was started with the sequential addition of 2 mM NAD^+^ and 60 units of lactate dehydrogenase. Absorbance at 340 nm was measured after 60 min using a Varioskan Flash plate reader.

### 2.5. Characterization of the cellular bioenergetics

An XF24 Extracellular Flux Analyzer (Agilent Technologies, Santa Clara, CA, USA) was used to determine the oxygen consumption rate (OCR) and the extracellular acidification rate (ECAR, a proxy for lactate formation) in the cell line A549. 12k cells were seeded in XF24-well microplates (Agilent Technologies) and transfected 24 hours later. Assays were carried out 48 hours after transfection. One hour prior to OCR and ECAR measurements, the culture medium was carefully removed, wells washed with 1 ml of assay medium and, finally, 500 μL of assay medium were added. The assay medium consisted of bicarbonate free DMEM medium pH 7.4 (Agilent Technologies, Santa Clara, CA, USA) supplemented with 2% of FBS, 2 mM glutamine and 5 mM glucose. Cells were maintained for 1 hour at 37°C in a CO_2_-free incubator before the start of the experiment.

The experimental protocol to determine the bioenergetics parameters was essentially as described in [21] which has been often termed as “mitochondrial stress test”. Basal OCR and ECAR were determined by performing four measurements and, subsequently, ATP turnover was assessed from the OCR decrease after the addition of oligomycin (1 μM). The maximal respiratory capacity and the spare respiratory capacity (SRC) were calculated from the OCR increase after two consecutive additions of the protonophore carbonyl cyanide p-(trifluoromethoxy)-phenylhydrazone (FCCP) after (0.6 and 0.4 μM). Finally, the respiratory chain inhibitors rotenone (1 μM) and antimycin A (1 μM) were added to determine the non-mitochondrial respiration.

ECAR values were corrected estimating the CO_2_ contribution to the ECAR signal from the correlation between the decreases in the OCR and the ECAR upon the addition of 100 mM 2-deoxyglucose (2-DOG). supplementary figures 1A & 1B show the steps followed to calculate the CO_2_ contribution: the difference in the ECAR value between the addition of 2-DOG and the inhibitors rotenone/antimycin was assigned as the CO_2_ contribution. The correlation between this ECAR value and the decrease in the OCR under the same conditions was used to estimate the CO_2_ contribution under a variety of experimental conditions (Suppl. Fig. 1C) that were compared with the lactate values determined enzymatically (Suppl. Fig. 1D). supplementary figure 1E shows that there exists a good correlation between the two experimental approaches (r_2_ = 0.827, p<0.002).

In all cases, at the end of each experiment, wells were washed with PBS and protein concentration determined by bicinchoninic acid assay using bovine serum albumin as standard.

### 2.6. Determination of reactive oxygen species

ROS levels were determined following the changes in fluorescence using the probe dihydroethidium (DHE). Sample analysis was carried out with an EPICS XL flow cytometer (Coulter, Hialeah, FL) collecting the fluorescence with a 620-nm band pass filter. Tripsinized cells were resuspended in standard growth medium and incubated for 1 hour at 37°C in the dark with 10 μM DHE.

### 2.7. Adenine nucleotides measurement

AMP, ADP and ATP levels were determined by reverse-phase high performance liquid chromatography (HPLC) essentially as described in [22]. 80k cells/well were seeded in p60 cell culture cells and transfected 24 hours later. Adenine nucleotide extraction was performed 48 hours after transfection. 30 min before the extraction, medium was replaced and cells incubated for 30 min at 37°C under the different experimental conditions. Nucleotides were quenched by the addition of 660 mM HClO_4_, 10 mM theophylline. Plates were quickly frozen using liquid nitrogen and subsequently stored at -80°C. Extracts were homogenized, centrifuged and supernatants containing the extracted nucleotides neutralized with 2.8 M K_3_PO_4_ and stored at -80°C overnight. Samples were thawed, centrifuged and supernatants were passed through a 0.45 μm filter. Analysis was performed using a Shimadzu Prominence chromatograph (Canby, Oregon, USA) using a C18 column (Mediterranea SEA18, Teknokroma, Spain). Peaks were identified according to the retention times of standard adenine nucleotides. Peak assignment was confirmed using samples treated with 2 μM oligomycin plus 50 mM 2-DOG in which the AMP and ADP peaks markedly increase. The cellular energy charge was calculated as indicated in [23].

### 2.8. Cell proliferation

Cell growth was monitored using a TC10 automated cell counter or by measuring the incorporation of nucleoside analog EdU (5-ethynyl-2’-deoxyuridine). For cell counting assays, 40k cells/well were seeded in p24 cell culture cells and transfected 24 hours later. Cell counting was performed 24 and 48 hours after transfection. The quantification of the nucleoside incorporation was performed with the Click-iT EdU microplate assay (Invitrogen, ThermoFisher Scientific, Lafayette, USA). 5k cells/well were seeded in 96-well cell culture plates and transfected 24 hours later. 48 hours after transfection, growth medium was replaced by fresh medium containing 20 μM EdU and mixed on a rotary shaker for 2 minutes. Cells were incubated for 4 hours at 37°C in a humidified 5% CO_2_ atmosphere and subsequently label incorporation was determined following manufacturer’s instructions. Amplex Red fluorescence was measured in a VarioskanFlash plate reader with the excitation at 568 nm and the emission at 585 nm.

### 2.9. Statistical analysis

All values are expressed as mean ± SEM. Differences between groups were determined using either two-tailed unpaired Student’s t-tests or the One-Way ANOVA test using the SigmaPlot software. Significant differences between groups are indicated as *p<0.05 or **p<0.01.

## 3. Results

### 3.1. OXPHOS and aerobic glycolysis

Silencing of UCP2 expression (supplementary figure 2) alters the cellular energy metabolism as determined from the analysis of the OCR and ECAR. The parameters calculated using the so-called “mitochondrial stress test” are summarized in figure 1. Significant differences were observed in the basal rate of respiration and in the ATP-turnover that was lower in the UCP2-silenced cells (Figs. 1A & 1B). The respiratory capacity (OCR in the presence of the uncoupler FCCP) was 40% higher than the basal rate (Fig. 1C). Although an apparent lower respiratory capacity in UCP2-silenced cells could be observed, it did not reach statistical significance (p=0.152). This result implies that mitochondrial mass remains almost unchanged. Interestingly, no differences were observed in the proton leakage rate (OCR in the presence of oligomycin, Fig. 1D). A change in the proton leak at high membrane potential would be expected if silencing was affecting the proton transport activity of an uncoupling protein [21].

**Fig 1.**
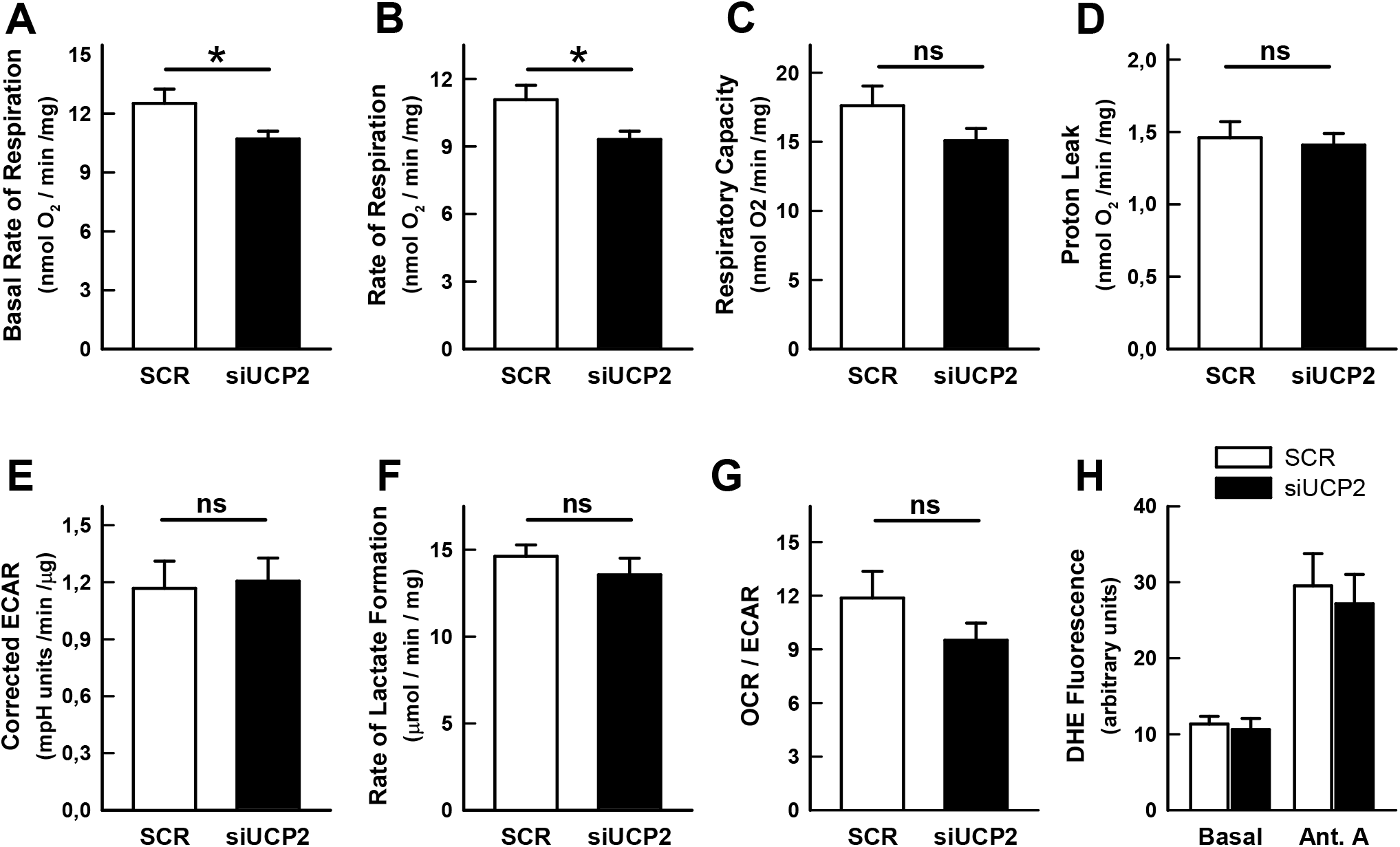
Effect of UCP2-silencing on the energy metabolism of the A549 cell line. A, Basal rate of respiration; B, Oligomycin sensitive respiration (ATP turnover); C, Uncoupled respiration (maximal respiratory capacity); D, Oligomycin-insensitive respiration (proton leakage rate); E, Basal rate of glycolysis corrected to take into account the CO_2_ contribution (see methods); F, Rate of lactate formation; G; OCR/ECAR ratio (bioenergetic profile); H, DHE fluorescence. In panels A-D and G, bars represent the mean ± SEM of 9 independent experiments with 3-5 technical replicates. Bars in panel F and H represent the mean ± SEM of 16 and 5 independent determinations respectively. Statistical significance: *p < 0.05, “ns” not significant difference.

In experiments with the Agilent-Seahorse Extracellular Flux Analyzer, the rate of aerobic glycolysis (lactate formation) can be estimated from ECAR values. Since the formation of bicarbonate from CO_2_ is also a net contributor to the extracellular acidification, to better ascertain the net rate of lactate formation, the CO_2_ contribution to the ECAR was estimated as indicated in the Methods Section (supplementary figure 1). The validity of this approach was confirmed by the enzymatic determination of the lactate formation using a variety of experimental conditions and comparing the values with ECAR determinations performed in parallel (supplementary figure 1E). The investigation of effect of UCP2 silencing on the rate of aerobic glycolysis, with the corrections mentioned above, revealed the absence of a significant increase in the ECAR signal (Fig. 1E) indicative of an unchanged rate of aerobic glycolysis and confirmed with the enzymatic determination of the concentration of lactate in the culture medium (Fig. 1F). The bioenergetic profile of the cells, as inferred from the ratio between the basal rates of respiration and glycolysis (OCR/ECAR ratio), would indicate that UCP2-silenced cells were slightly more glycolytic although the difference did not reach statistical significance (p=0.202) (Fig. 1G).

### 3.2. Reactive oxygen species

The activity of the UCPs has been proposed to modulate the formation of mitochondrial ROS. If UCP2 silencing lowers the rate of respiration, an increase in mitochondrial ROS formation could be expected. Figure 1H shows that silencing does not change ROS levels as determined with the fluorescent probe DHE. Results with the complex III inhibitor antimycin A, used as a control, demonstrate that under our experimental conditions it is possible to detect increases in ROS levels upon inhibition of respiration.

### 3.3. Adenine nucleotide levels

The combination of a lower rate of respiration without a compensatory increase in aerobic glycolysis reflects a lower energy metabolism metabolic activity in the silenced cells that could be the result of three possible situations: (i) an increased metabolic efficiency, (ii) a lower cellular energy demand or (iii) a metabolic failure that compromises the cellular energy levels. Figure 2 presents the results of the analysis by HPLC of the cellular content of adenine nucleotides levels and shows that UCP2-silenced cells presented un unchanged cellular energy charge and thus with no differences in the ADP/ATP or the AMP/ATP ratios. These results exclude the possibility of a metabolic failure. It is interesting to note that, in the two cell types, the adenine nucleotides pools responded identically to the inhibition of the metabolic pathways that sustain the energy levels.

**Fig. 2.**
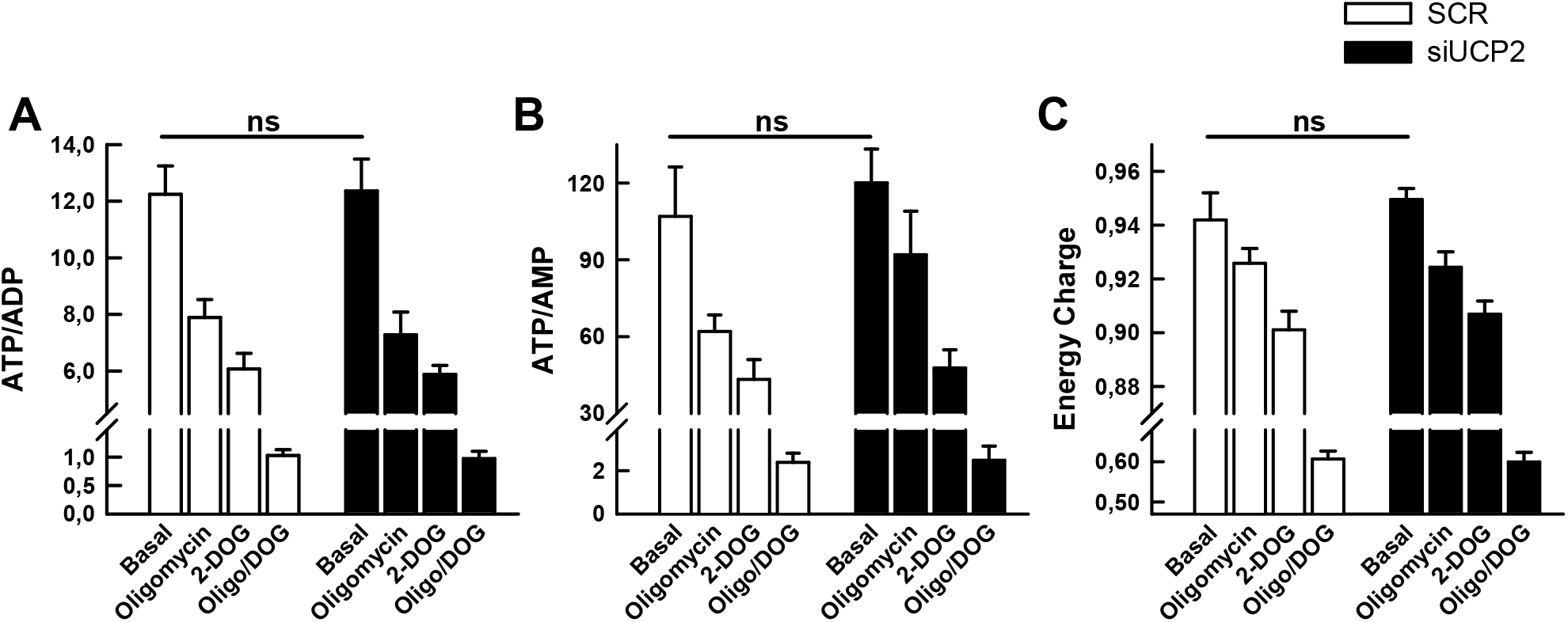
Effect of UCP2 silencing on the adenine nucleotide levels of the A549 cell line. A, ATP/ADP ratio; B, ATP/AMP ratio; C, Energy charge as defined in [23]. Nucleotide extracts were prepared con control A549 cells (SCR) and UCP2-silenced cells (siUCP2) either untreated (“Basal”) or treated for 30 min with 1 μM oligomycin, 100 mM 2-DOG or 1 μM oligomycin plus 100 mM 2-DOG (Oligo/DOG). Bars represent the mean ± SEM of 10-14 independent experiments. “ns” denotes the absence of significant differences between “Basal” values.

### 3.4. Cell growth

Since UCP2 has been involved in the Warburg effect which occurs in proliferative cells there was the possibility that the decreased energy demand was due to a lower proliferation. Figure 3 shows the effect of UCP2 silencing in the proliferation of the A549 cell line. Results show a significant decrease in the number of cells 48 hours after transfection (p=0.003). The rate of incorporation of the nucleoside analog EdU over a period of 4 hours (Fig. 3B) followed a trend consistent with the decrease in cell number although it did not reach statistical significance (p= 0.114).

**Fig. 3.**
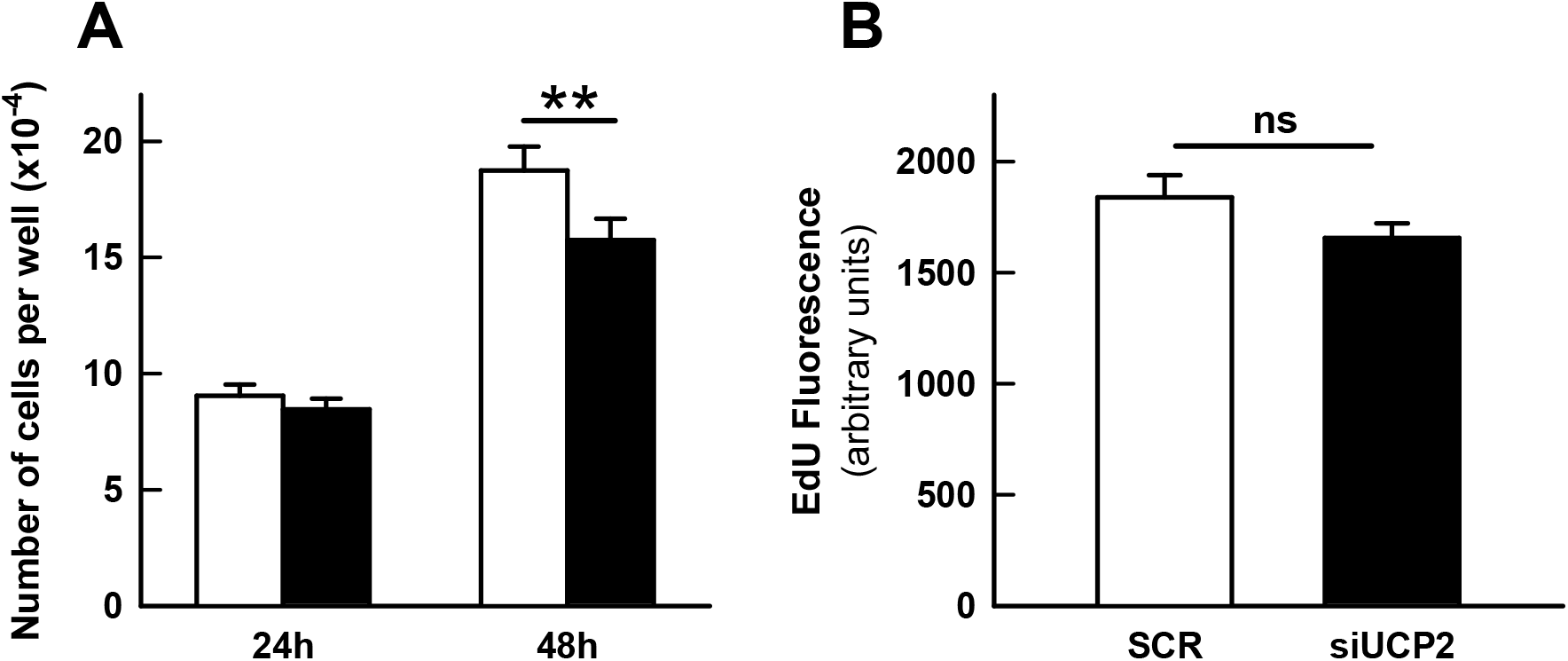
Effect of UCP2 silencing in the proliferation of the A549 cell line. A, Number of cells per well determined 24 and 48 hours after transfection. Bars represent the mean ± SEM of 4 independent experiments with 3 technical replicates in each experiment. B, Incorporation of the fluorescent nucleoside analog EdU over a period of 4 hours, 48 hours after transfection. Bars represent the mean ± SEM of 4 independent experiments with 12-16 technical replicates in each experiment. Statistical significance: **p < 0.01, “ns” not significant difference.

### 3.5. Chromane derivatives

We have previously described a set of chromane derivatives that inhibit the proton conductance of yeast mitochondria expressing UCP1 [24]. We also showed that, in HT-29 colon carcinoma cells, the chromane CSIC-E379 inhibits mitochondrial respiration, causing oxidative stress and decreasing their viability. We have reviewed the effects of the chromane derivative CSIC-E379 in our new cellular model where we can test the effect in UCP2-silenced cells. Figure 4 Results showed that the chromane inhibits cellular respiration even in the absence of UCP2 (Fig. 4A) and the inhibition led to (i) an increase in ROS formation (Fig. 4B) and (ii) an increase in lactate formation (Fig. 4C). These two combined effects are indicative of an unspecific inhibitory effect of the chromane on the mitochondrial respiratory chain. These toxic effects where not detected during the screening experiments that led to the identification of these novel inhibitors of UCP1 and UCP2 [24]. Thus the chromane CSIC-E379 had no effects on mitochondria isolated from. *cerevisiae* that did not express either UCP1 or UCP2. Since these mitochondria do not possess a proton-translocating rotenone-sensitive NADH dehydrogenase, we assume that the observed respiratory inhibition must be due to the interaction of the chromane with the mammalian complex I.

**Fig. 4.**
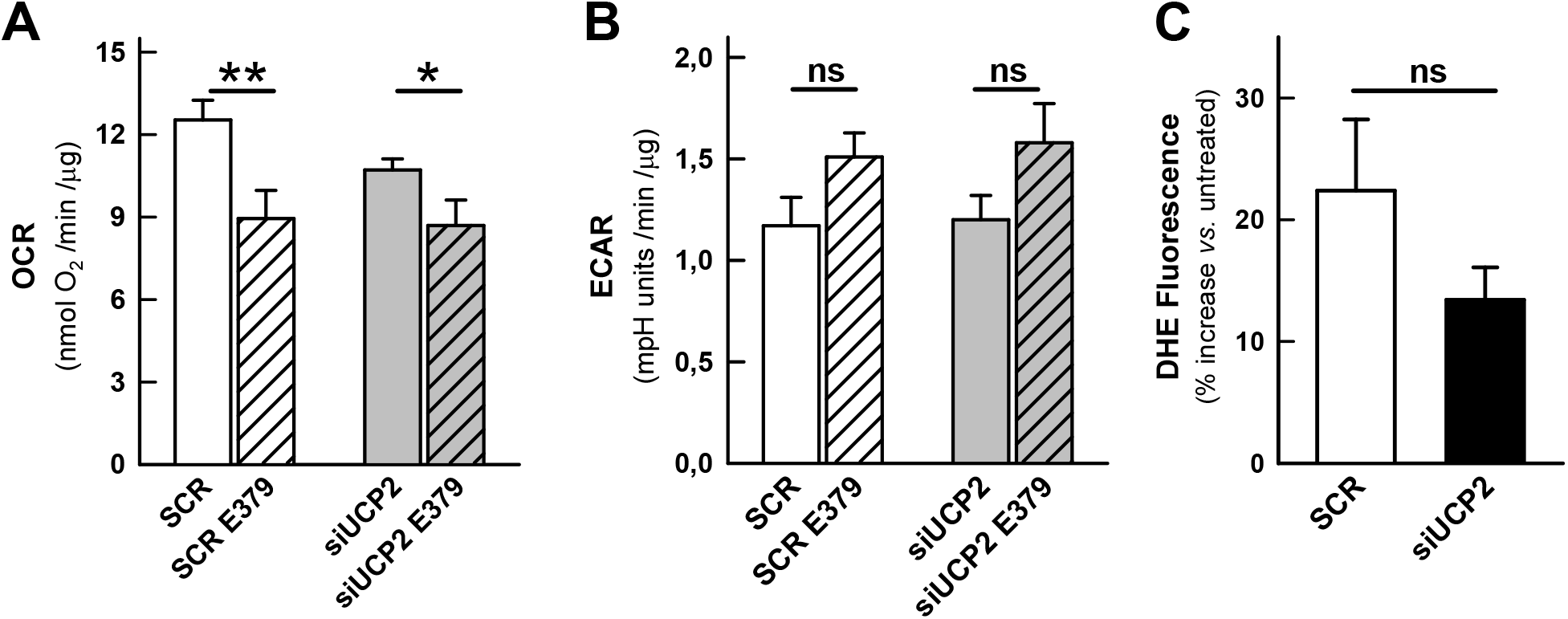
Effect of the chromane derivative CSIC-E379 in control and UCP2-silenced cells. (A) Basal rate of respiration and (B) Basal rate of glycolysis of control cells (SCR) and UCP2-silenced cells treated for 24 hours with 20 μM CSIC-E379. Bars represent the mean ± SEM of 6-8 independent experiments with 3-5 technical replicates. C, DHE fluorescence increase in control and UCP2-silenced cells after cell treatment for 24 hours with 20 μM CSIC-E379 of 5 independent experiments. Values are expressed as the percentage of fluorescent increase with respect to untreated cells. Statistical significance: *p < 0.05, **p < 0.01, “ns” not significant difference.

## 4. Discussion

UCP2 was discovered in 1997 due to its sequence homology to UCP1 [25]. It was found that, while UCP1 was only expressed in brown adipose tissue, UCP2 had a wide tissue distribution [26]. The gene localization in chromosomal regions linked to hyperinsulinaemia and obesity suggested that UCP2 could have a role in the control of energy balance and, additionally, in the response to inflammatory stimuli [25]. The publication, that same year, of the leptin induction of UCP2 in white adipose tissue together with enzymes involved in fatty acid oxidation further supported a thermogenic role [27]. The involvement of UCP2 in the control of mitochondrial ROS formation was soon uncovered [28] and, subsequently, reports have continued to appear pointing to UCP2 as an inducible protein involve in the defense against oxidative stress [4,7,12,13,20,29,30]. The mechanistic basis would lie on the steep dependence of the mitochondrial formation of superoxide with membrane potential (Figure 5), thus, “mild” respiratory uncoupling or the state 4 to state 3 transition would be sufficient to lower the membrane potential 10-20 mV and drastically reduce ROS formation [31-35] and figure 5. These findings implied that, like UCP1, UCP2 was a carrier that would uncouple OXPHOS by increasing proton leakage. However, UCP2-mediated proton permeation has been an elusive biochemical activity which remains controversial [6,8,10,11,19]. Under our experimental conditions, UCP2 silencing did not lead to differences in proton leakage, as determined from the rate of respiration in the presence of oligomycin (Figure 1C). Similar results have been reported with isolated mitochondria from UCP2 knockout mice [36-37] and with brain mitochondria from transgenic mice overexpressing UCP2 [38]. The dependence of the proton leak with membrane potential [31,35] rules out the possibility of a significant UCP2-mediated uncoupling activity when mitochondria are actively phosphorylating ADP.

**Fig. 5.**
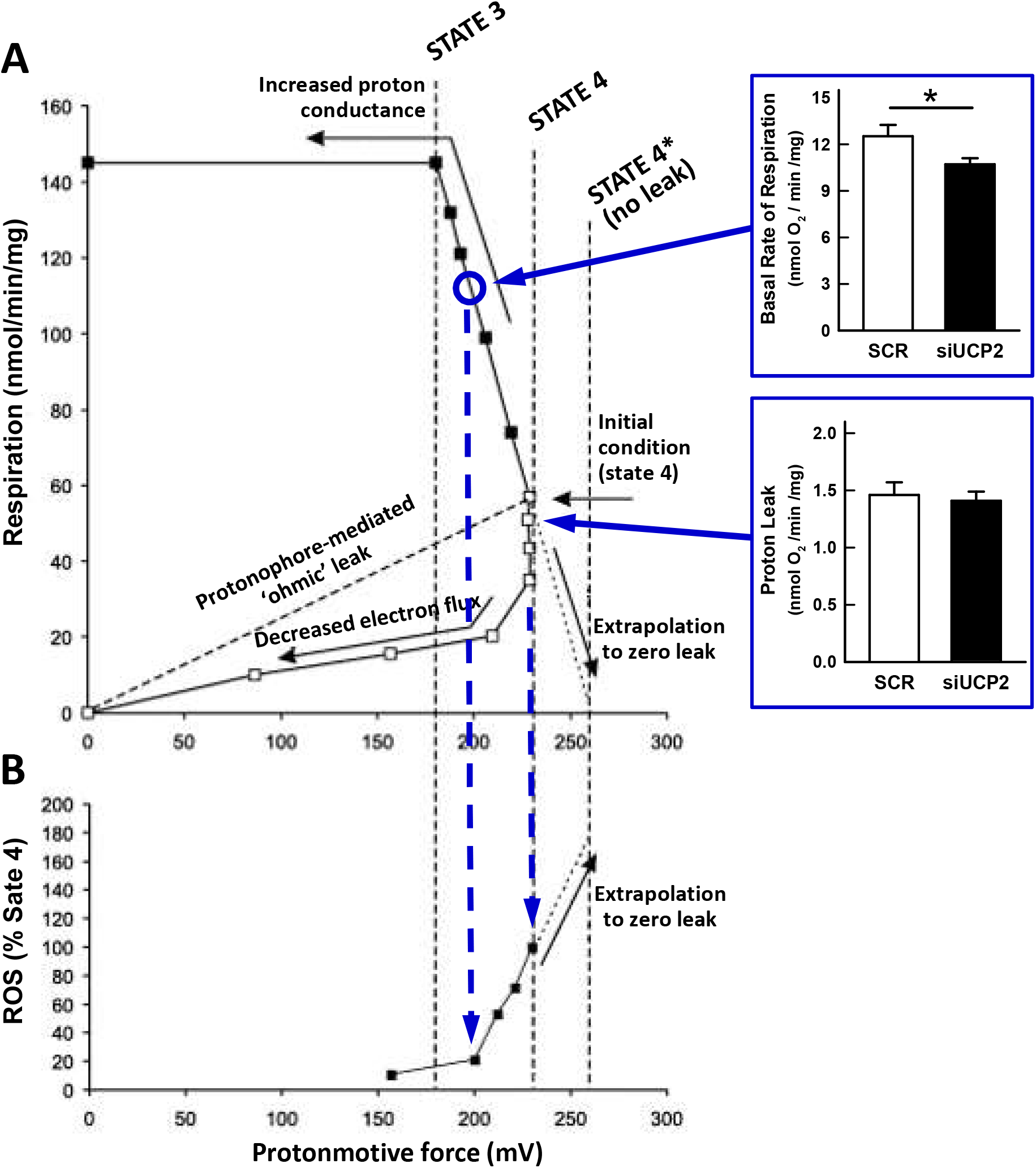
Experimental data relating mitochondrial respiration, protonmotive force and ROS production (reproduced with permission from [35]). (A) Relationships between respiration and protonmotive force for brown fat mitochondria that are either exposed to an increasing demand on the proton current (uncoupler titration, closed symbols) or whose substrate supply is progressively restricted (substrate titration, open symbols). Original data from [48,49]. (B) Data replotted from [31] showing the relationship between H_2_O_2_ generation and protonmotive force. See [35] for further details. Histograms included are taken from figure 1 and graphically show how the observed state 3 rates (blue circle would correspond to 70% of the uncoupled rate) should correspond to low rates of ROS formation (dashed arrows).

The presence of UCP2 in tumor cells was another early finding [39] and is providing important clues to its biochemical function. The discovery that drug-resistant tumor cells express high levels of UCP2 [13,40], that was linked to protection against oxidative stress, pointed to UCP2 as a new drug target for cancer treatment [14-16,24,41]. In addition, a growing number of reports implicated UCP2 in the metabolic adaptations of cancer cells, the so-called Warburg effect [10,14,17-19,42-44]. However, a clue to this function could lie in the strong regulation of UCP2 translation by glutamine [6,9,45], a key substrate required for cell proliferation [46-47]. Thus, glutaminolysis provides anaplerotic precursors that drive a glucose-independent Krebs cycle in which intermediates like citrate or oxaloacetate will be exported to be used in the synthesis of macromolecules. In line with these concepts, the discovery that UCP2 is a C4-metabolite carrier that facilitates the export of malate, oxaloacetate and aspartate from the mitochondrial matrix seems to provide a mechanistic explanation for the role of UCP2 in the metabolic adaptations of cancer cells [9]. In the present work we have observed that UCP2 silencing decreases the proliferation of the cell line A549 (Figure 3) consistent with the proposed involvement of UCP2 in the Warburg effect. Our data suggest that this lower growth could be the cause of the observed decrease in mitochondrial respiration, that is not compensated by an increased in glycolysis (Figure 1) and that does not lead to a decrease in the cellular energy charge (Figure 2). In other words, these results are indicative of a decrease in the cellular ATP demand as a result of the decreased cell proliferation.

Our study also provides clues to the role of UCP2 in the control of mitochondrial ROS formation. Under our experimental conditions, the effect of UCP2 silencing is to decrease the basal rate of respiration, i.e. the state 3 rate (Figure 1A). It is important to note the magnitude of the inhibition of basal respiration by oligomycin in control cells (88,3% ± 0,7) and that the FCCP uncoupled rate is only 40% higher than the basal rate, implying that mitochondria are actively synthesizing ATP and, therefore, ought to be partially depolarized. The relationship between the rate of respiration, membrane potential and rate of ROS formation presented in Figure 5 (reproduced with permission from [35]) explicitly portraits that, under those conditions, if UCP2 were to catalyze proton leakage it would have no influence on the rate of ROS formation. Accordingly, no differences were observed in DHE fluorescence (Figure 1H). It has already been pointed out that the UCP2-mediated export of matrix C4 metabolites could lower redox pressure on the mitochondrial respiratory that could result in a lower ROS production [9] although this would be barely detectable when mitochondria are actively phosphorylating ADP.

As we have previously stated, UCP2 is upregulated in many physiological situations where there is oxidative stress and that its upregulation has also been linked to the Warburg effect. Proliferating cells, and cancer cells in particular, need metabolic intermediates, reducing power (NADPH) and ATP to synthesize biomass. Additionally, cancer cells have to cope with a high intrinsic oxidative stress and the pentose phosphate pathway is one of the routes that regenerates the NADPH required for ROS detoxification. The increased uptake of glucose and glutamine should serve all these purposes and cellular metabolism ought to be adjusted accordingly [50,51]. Interestingly, it has already been shown that glutamine metabolism supports NADPH production, mainly through the cytosolic malic enzyme [52], and also the generation of biosynthetic intermediates. UCP2 would appear to play a key role in these two processes through the export of mitochondrial malate, oxaloacetate and aspartate [9]. In this context, UCP2 could indeed be considered as an element of the cellular antioxidant defense.

## Abbreviations

DHE,: dihydroethidium;
ECAR,: extracellular acidification rate;
FBS,: fetal bovine serum;
FCCP,: Carbonyl cyanide p-(trifluoromethoxy)-phenylhydrazone;
OCR,: oxygen consumption rate;
OXPHOS: oxidative phosphorylation;
ROS,: reactive oxygen species;
UCP,: uncoupling protein.

## Acknowledgements

We thank Dr. Juan P. Bolaños for critical comments on the manuscript. The work has been funded by grants from the Spanish Ministry of Science and Innovation (SAF2010-20256, PID2019-108166GB-I00 and Redox Biology and Medicine Network RED2018-102576-T).

## Author contributions

**Jessica Segalés:** Methodology, Investigation, Formal analysis. **Carlos Sánchez-Martín, Aleida Pujol** and **Marta Martín-Ruiz**: Investigation, Formal analysis. **Eduardo Rial:** Conceptualization, Funding acquisition, Methodology, Formal analysis, Writing-Review Editing. All the authors approved the final version of the manuscript.

## References

[1] J.J. Ruprecht, E.R.S. Kunji, Structural mechanism of transport of mitochondrial carriers, Annu Rev Biochem 90 (2021) 535–558.

[2] D.G. Nicholls, R.M. Locke, Thermogenic mechanisms in brown fat, Physiol Rev 64 (1984) 1–64.

[3] F. Bouillaud, E. Couplan, C. Pecqueur, D. Ricquier, Homologues of the uncoupling protein from brown adipose tissue (UCP1): UCP2, UCP3, BMCP1 and UCP4, Biochim Biophys Acta 1504 (2001) 107–19.

[4] M.D. Brand, C. Affourtit, T.C. Esteves, K. Green, A.J. Lambert, S. Miwa, J.L. Pakay, N. Parker, Mitochondrial superoxide: production, biological effects, and activation of uncoupling proteins, Free Radic Biol Med 37 (2004), 755–767.

[5] B. Cannon, I.G. Shabalina, T.V. Kramarova, N. Petrovic N, J. Nedergaard, Uncoupling proteins: a role in protection against reactive oxygen species—or not?, Biochim Biophys Acta 1757 (2006) 449–458.

[6] F. Bouillaud, UCP2, not a physiologically relevant uncoupler but a glucose sparing switch impacting ROS production and glucose sensing, Biochim Biophys Acta 2009 (1787), 377–383

[7] R.J. Mailloux, M.E. Harper, Uncoupling proteins and the control of mitochondrial reactive oxygen species production, Free Radic Biol Med 51 (2011) 1106–1115

[8] I.G. Shabalina, J. Nedergaard, Mitochondrial (‘mild’) uncoupling and ROS production: physiologically relevant or not?, Biochem Soc Trans 39 (2011) 1305–1309

[9] A. Vozza, G. Parisi, F. De Leonardis, F.M. Lasorsa, A. Castegna, D. Amorese, R. Marmo, V.M. Calcagnile, L. Palmieri, D. Ricquier, E. Paradies, P. Scarcia, F. Palmieri, F. Bouillaud, G. Fiermonte, UCP2 transports C4 metabolites out of mitochondria, regulating glucose and glutamine oxidation, Proc Natl Acad Sci U S A 111 (2014) 960–965

[10] F. Bouillaud, M.C. Alves-Guerra, D. Ricquier, UCPs, at the interface between bioenergetics and metabolism, Biochim Biophys Acta 1863 (2016) 2443–56.

[11] D.G. Nicholls, Mitochondrial proton leaks and uncoupling proteins, Biochim Biophys Acta Bioenerg 1862 (2021) 148428.

[12] J. Hirschenson, E. Melgar-Bermudez, R.J. Mailloux, The uncoupling proteins: a systematic review on the mechanism used in the prevention of oxidative stress, Antioxidants 11 (2022) 322.

[13] M.E. Harper, A. Antoniou, E. Villalobos-Menuey, A. Russo, R. Trauger, M. Vendemelio, A. George, R. Bartholomew, D. Carlo, A. Shaikh, J. Kupperman, E.W. Newell, I.A. Bespalov, S.S. Wallace, Y. Liu, J.R. Rogers, G.L. Gibbs, J.L. Leahy, R.E. Camley, R. Melamede, M.K. Newell, Characterization of a novel metabolic strategy used by drug-resistant tumor cells, FASEB J 16 (2002) 1550–1557.

[14] I. Samudio, M. Fiegl, M. Andreeff, Mitochondrial uncoupling and the Warburg effect: molecular basis for the reprogramming of cancer cell metabolism, Cancer Res 69 (2009) 2163–6.

[15] G. Baffy, Uncoupling protein-2 and cancer, Mitochondrion 10 (2010) 243–252

[16] G. Baffy, Z. Derdak, S.C. Robson, Mitochondrial recoupling: a novel therapeutic strategy for cancer?, Br J Cancer 105 (2011) 469–474

[17] P. Esteves, C. Pecqueur, C. Ransy, C. Esnous, V. Lenoir, F. Bouillaud, A.L. Bulteau, A. Lombès, C. Prip-Buus, D. Ricquier, M.C. Alves-Guerra, Mitochondrial retrograde signaling mediated by UCP2 inhibits cancer cell proliferation and tumorigenesis, Cancer Res 74 (2014) 3971–3982.

[18] A. Sreedhar, P. Petruska, S. Miriyala, M. Panchatcharam, Y. Zhao, UCP2 overexpression enhanced glycolysis via activation of PFKFB2 during skin cell transformation, Oncotarget 8 (2017) 95504–95515.

[19] A. Rupprecht, R. Moldzio, B. Mödl, E.E. Pohl, Glutamine regulates mitochondrial uncoupling protein 2 to promote glutaminolysis in neuroblastoma cells, Biochim Biophys Acta 1860 (2019) 391–401.

[20] A. Anedda, E. Rial, M.M. González-Barroso, Metformin induces oxidative stress in white adipocytes and raises uncoupling protein 2 levels, J Endocrinol 199 (2008) 33–40.

[21] M.D. Brand, D.G. Nicholls, Assessing mitochondrial dysfunction in cells, Biochem J 435 (2011) 297–312.

[22] D. De Korte, W.A. Haverkort, A.H. van Gennip, D. Roos, Nucleotide profiles of normal human blood cells determined by high-performance liquid chromatography, Anal Biochem 147 (1985) 197–209.

[23] D.E. Atkinson, The energy charge of the adenylate pool as a regulatory parameter. Interaction with feedback modifiers, Biochemistry 7 (1968) 4030–4034.

[24] E. Rial, L. Rodríguez-Sánchez, P. Aller, A. Guisado, M.M. González-Barroso, E. Gallardo-Vara, M. Redondo-Horcajo, E. Castellanos, R. Fernández de la Pradilla, A. Viso, Development of chromanes as novel inhibitors of the uncoupling proteins, Chem Biol 18 (2011) 264–274.

[25] C. Fleury, M. Neverova M, S. Collins, S. Raimbault, O. Champigny, C. Levi-Meyrueis, F. Bouillaud, M.F. Seldin, R.S. Surwit, D. Ricquier, C.H. Warden, Uncoupling protein-2: a novel gene linked to obesity and hyperinsulinemia, Nat Genet 15 (1997) 269–272.

[26] D. Ricquier, F. Bouillaud, The uncoupling protein homologues: UCP1, UCP2, UCP3, StUCP and AtUCP, Biochem J 345 (2000) 161–179.

[27] Y.T. Zhou YT M. Shimabukuro, K. Koyama, Y. Lee, M.Y. Wang, F. Trieu, C.B. Newgard, R.H. Unger, Induction by leptin of uncoupling protein-2 and enzymes of fatty acid oxidation, Proc Natl Acad Sci U S A 94 (1997) 6386–6390.

[28] A. Nègre-Salvayre, C. Hirtz C, G. Carrera, R. Cazenave, M. Troly, R. Salvayre, L. Pénicaud, L. Casteilla, A role for uncoupling protein-2 as a regulator of mitochondrial hydrogen peroxide generation, FASEB J 11 (1997) 809–815.

[29] M.D. Brand, T.C. Esteves, Physiological functions of the mitochondrial uncoupling proteins UCP2 and UCP3, Cell Metab 2 (2005) 85–93.

[30] S. Cadenas, Mitochondrial uncoupling, ROS generation and cardioprotection, Biochim Biophys Acta Bioenerg 1859 (2018) 940–950.

[31] S.S. Korshunov, V.P. Skulachev, A.A. Starkov, High protonic potential actuates a mechanism of production of reactive oxygen species in mitochondria, FEBS Lett 416 (1997) 15–18.

[32] V.P. Skulachev, Role of uncoupled and non-coupled oxidations in maintenance of safely low levels of oxygen and its one-electron reductants, Q Rev Biophys 29 (1996) 169–202.

[33] S. Miwa, M.D. Brand, Mitochondrial matrix reactive oxygen species production is very sensitive to mild uncoupling, Biochem Soc Trans 31 (2003) 1300–1301.

[34] A.A. Starkov, G. Fiskum, Regulation of brain mitochondrial H2O2 production by membrane potential and NAD(P)H redox state, J Neurochem 86 (2003) 1101–1107.

[35] D.G. Nicholls, Mitochondrial membrane potential and aging, Aging Cell 3 (2004) 35–40.

[36] E. Couplan, M.M. Gonzalez-Barroso, M.C. Alves-Guerra, D. Ricquier, M. Goubern, F. Bouillaud, No evidence for a basal, retinoic, or superoxide-induced uncoupling activity of the uncoupling protein 2 present in spleen or lung mitochondria, J Biol Chem 277 (2002) 26268– 26275.

[37] A. Kukat, S.A. Dogan, D. Edgar, A. Mourier, C. Jacoby, P. Maiti, J. Mauer, C. Becker, K. Senft, R. Wibom, A.P. Kudin, K. Hultenby, U. Flögel, S. Rosenkranz, D. Ricquier, W.S. Kunz, A. Trifunovic, Loss of UCP2 attenuates mitochondrial dysfunction without altering ROS production and uncoupling activity, PLoS Genet 10 (2014) e1004385.

[38] G. Mattiasson, M. Shamloo, G. Gido, K. Mathi, G. Tomasevic, S. Yi, C.H. Warden, R.F. Castilho, T. Melcher, M. Gonzalez-Zulueta, K. Nikolich, T. Wieloch, Uncoupling protein-2 prevents neuronal death and diminishes brain dysfunction after stroke and brain trauma. Nature Med 9 (2003) 1062–1068.

[39] M.V. Carretero, L. Torres, U. Latasa, E.R. García-Trevijano, J. Prieto, J.M. Mato, M.A. Avila, Transformed but not normal hepatocytes express UCP2, FEBS Lett 439 (1998) 55–58.

[40] Z. Derdak, N.M. Mark, G. Beldi, S.C. Robson, J.R. Wands, G. Baffy, The mitochondrial uncoupling protein-2 promotes chemoresistance in cancer cells, Cancer Res 68 (2008) 2813–2819.

[41] P.G. Pons, M. Nadal-Serrano, M. Torrens-Mas, A. Valle, J. Oliver, P. Roca, UCP2 inhibition sensitizes breast cancer cells to therapeutic agents by increasing oxidative stress, Free Radic Biol Med 86 (2015) 67–77.

[42] Brandi J, Cecconi D, Cordani M, Torrens-Mas M, Pacchiana R, Dalla Pozza E, Butera G, Manfredi M, Marengo E, Oliver J, Roca P, Dando I, Donadelli M, The antioxidant uncoupling protein 2 stimulates hnRNPA2/B1, GLUT1 and PKM2 expression and sensitizes pancreas cancer cells to glycolysis inhibition, Free Radic Biol Med 101 (2016) 305–316.

[43] J. Zhang, I. Khvorostov, J.S. Hong, Y. Oktay, L. Vergnes, E. Nuebel, P.N. Wahjudi, K. Setoguchi, G. Wang, A. Do, J.H. Jung, J.M. McCaffery, I.J. Kurland, K. Reue, W.N. Lee, C.M. Koehler, M.A. Teitell, UCP2 regulates energy metabolism and differentiation potential of human pluripotent stem cells, EMBO J 30 (2011) 4860–4873.

[44] S. Raho, L. Capobianco, R. Malivindi, A. Vozza, C. Piazzolla, F. De Leonardis, R. Gorgoglione, P. Scarcia, F. Pezzuto, G. Agrimi, S.N. Barile, I. Pisano, S.J. Reshkin, M.R. Greco, R.A. Cardone, V. Rago, Y. Li, C.M.T. Marobbio, W. Sommergruber, C.L. Riley, F.M. Lasorsa, E. Mills, M.C. Vegliante, G.E. De Benedetto, D. Fratantonio, L. Palmieri, V. Dolce, G. Fiermonte, KRAS-regulated glutamine metabolism requires UCP2-mediated aspartate transport to support pancreatic cancer growth, Nature Metab 2 (2020) 1373–1381.

[45] C. Hurtaud, C. Gelly, Z. Chen, C. Lévi-Meyrueis, F. Bouillaud, Glutamine stimulates translation of uncoupling protein 2 mRNA, Cell Mol Life Sci 64 (2007) 1853–1860.

[46] B.J. Altman, Z.E. Stine, C.V Dang, From Krebs to clinic: glutamine metabolism to cancer therapy, Nat Rev Cancer 16 (2016) 619–634.

[47] M.G. Vander Heiden, R.J. DeBerardinis, Understanding the intersections between metabolism and cancer biology, Cell 168 (2017) 657–669.

[48] D.G. Nicholls, The influence of respiration and ATP hydrolysis on the proton electrochemical potential gradient across the inner membrane of rat liver mitochondria as determined by ion distribution, Eur J Biochem 50 (1974) 305–315.

[49] D.G. Nicholls, The effective proton conductances of the inner membrane of mitochondria from brown adipose tissue: dependency on proton electrochemical gradient, Eur J Biochem 77 (1977) 349–356.

[50] X. Chen, Y. Qian, S. Wu, The Warburg effect: evolving interpretations of an established concept, Free Radic Biol Med 79 (2015) 253–263.

[51] A.C. Boese, S. Kang, Mitochondrial metabolism-mediated redox regulation in cancer progression, Redox Biol 42 (2021) 101870.

[52] R.J. DeBerardinis, A. Mancuso, E. Daikhin, I. Nissim, M. Yudkoff, S. Wehrli, C.B. Thompson, Beyond aerobic glycolysis: transformed cells can engage in glutamine metabolism that exceeds the requirement for protein and nucleotide synthesis, Proc Natl Acad Sci U S A 104 (2007) 19345–19350.

