## Supplemental Figures for "Role of UCP2 in the energy metabolism of the cancer cell line A549"

**LEGENDS TO SUPPLEMENTARY FIGURES**

Supplementary Fig. 1. Experimental approach to correct ECAR data and remove the $CO_2$ contribution to the acidification and thus better estimate the rate of lactate formation.
A, OCR measurement in A549 cells with the sequential additions of 100 mM 2-DOG, 1 mM oligomycin and 1 μM rotenone plus 1 μM antimycin A. For subsequent calculations, the relevant change in OCR is indicated with the blue double arrow. B, ECAR determination following the same experimental protocol than in A. The ECAR variations that should be due either to lactate or to bicarbonate are indicated. C, Variation in the ECAR signal after the subtraction of the calculated $CO_2$ contribution with cells under experimental conditions that should lead to changes in lactate formation. D, Rate of lactate formation determined enzymatically as in C. Experimental conditions for data in panels C & D are the following: "Basal", untreated A549 cells; "2-DOG 30", A549 cells after treatment with 30 mM 3-DOG; "2-DOG 100", A549 cells after treatment with 100 mM 2-DOG; "SCR CTRL", untreated control A549 transfected cells; "SCR METF" control A549 transfected cells treated for 1 hour with 1 mM metformin; "siUCP2 CTRL", UCP2-silenced A549 untreated; "siUCP2 METF" UCP2-silenced A549 treated for 1 hour with 1 mM metformin; "Oligomycin", A549 cells treated with 1 μM oligomycin; "Rot/AA +2DOG", A549 cells treated with 100 mM 2-DOG,1 μM rotenone and 1 μM antimycin A. E, Correlation between the values presented in histograms C & D ($r^2 = 0.827$, $p<0.002$).

Supplementary Fig. 2. A & B, representative Western blots showing the effect of UCP2-silencing on the expression levels of the protein. "Spleen Mit." are mitochondria isolated from mouse spleen used as markers due to the high UCP2 content; A549, cell extracts from untransfected A549 cells; SCR, cell extracts from control A549 cell transfected with scrambled siRNA. siUCP2, cell extracts from UCP2-silenced cells. Luminescence signal is saturated in some of the lanes to ensure that low UCP2 protein levels can be detected. All extracts are from independent cell preparations.

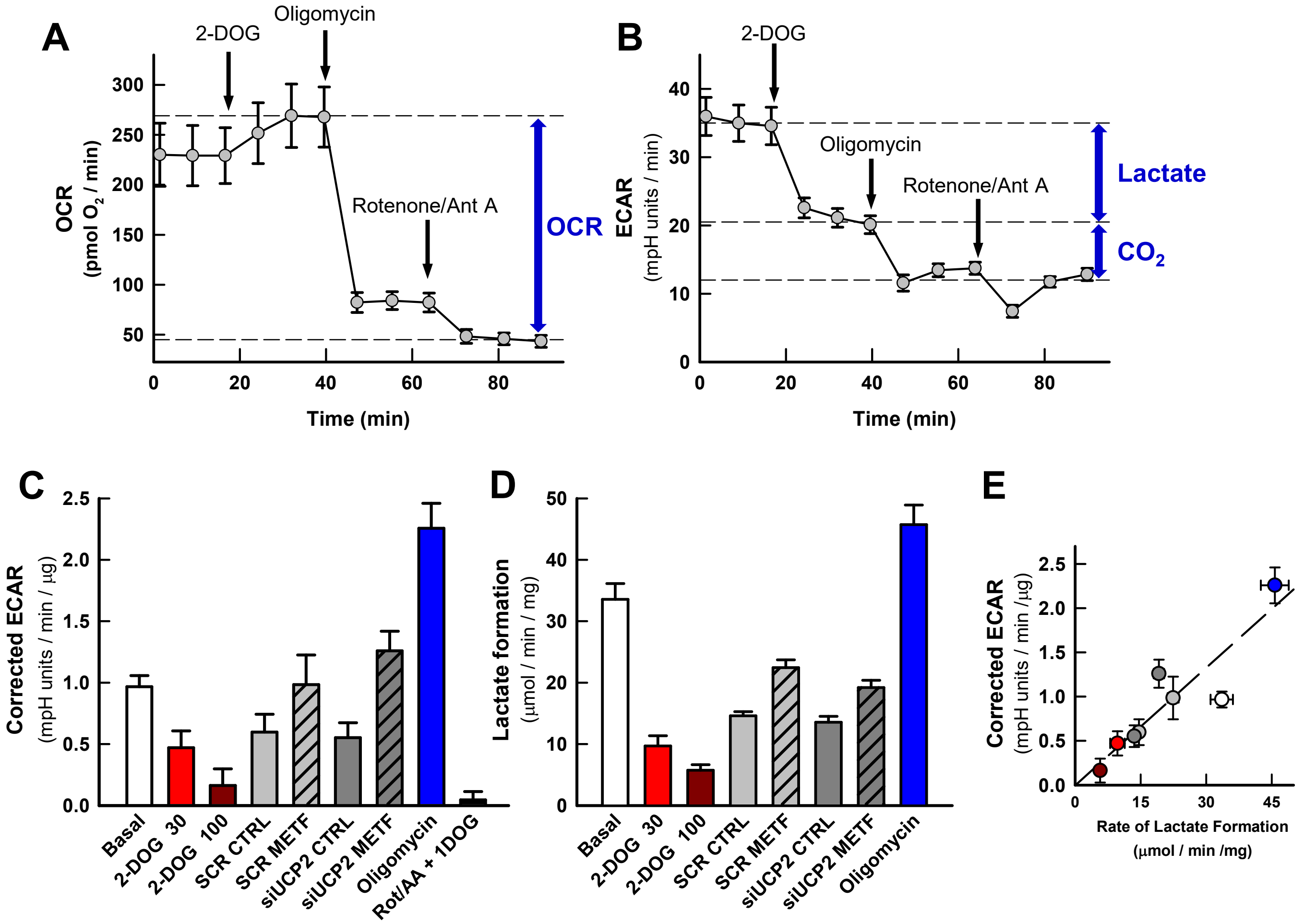


**Supplementary Figure 1**

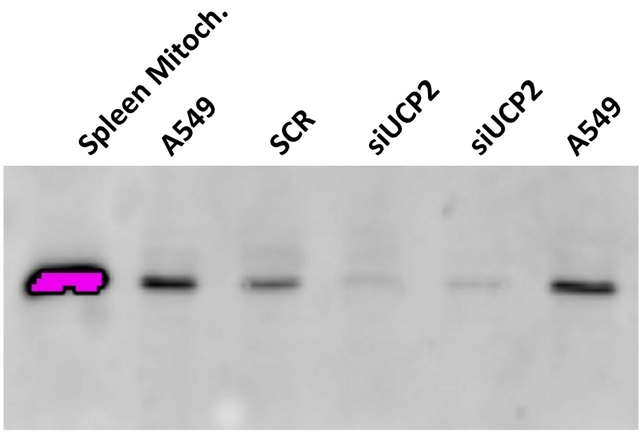


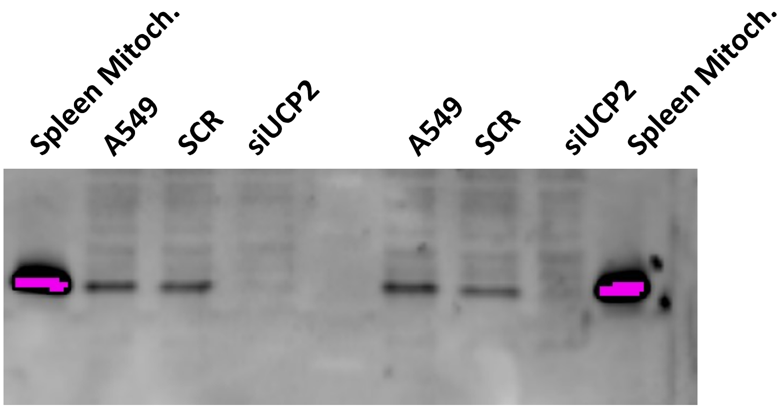


**Supplementary Figure 2**
